# Polo-like kinase 1 independently controls microtubule-nucleating capacity and size of the centrosome

**DOI:** 10.1101/2020.10.07.328740

**Authors:** Midori Ohta, Zhiling Zhao, Di Wu, Shaohe Wang, Jennifer L. Harrison, J. Sebastián Gómez-Cavazos, Arshad Desai, Karen Oegema

## Abstract

Centrosomes are composed of a centriolar core surrounded by a pericentriolar material (PCM) matrix that docks microtubule-nucleating γ-tubulin complexes. During mitotic entry, the PCM matrix increases in size and nucleating capacity in a process called centrosome maturation. Polo-like kinase 1 (PLK1) localizes to centrosomes and phosphorylates PCM matrix proteins to drive their self-assembly, which leads to PCM expansion; this expansion has been assumed to passively increase microtubule nucleation to support spindle assembly. Here, we show that PLK1 directly controls the generation of binding sites for γ-tubulin complexes on the PCM matrix, independently of PCM expansion. Selective inhibition of PLK1-dependent γ-tubulin docking leads to spindle defects and impaired chromosome segregation, without affecting PCM expansion, highlighting the importance of phospho-regulated centrosomal γ-tubulin docking sites in spindle assembly. Inhibiting both γ-tubulin docking and PCM expansion by mutating substrate target sites fully accounts for the actions of PLK-1 in transforming the centrosome during mitotic entry.

**Summary Statement:** Polo-like kinase 1-mediated physical expansion of centrosomes during mitotic entry is proposed to passively increase their microtubule nucleating capacity. Ohta et al. show instead that generation of microtubule-nucleating sites is directly controlled by Polo-like kinase 1, independently of centrosome size.

## INTRODUCTION

During cell division in metazoans, centrosomes catalyze microtubule generation for assembly of the mitotic spindle (Basto et al., 2006; Bazzi and Anderson, 2014; Khodjakov and Rieder, 2001; Meitinger et al., 2016; Pintard and Bowerman, 2019; Sir et al., 2013; Wong et al., 2015). Centrosomes comprise a centriolar core surrounded by an extended pericentriolar material (PCM) matrix that nucleates and anchors microtubules (Kellogg et al., 1994; Mennella et al., 2014; Woodruff et al., 2014). The PCM matrix forms via the self-assembly of large coiled-coil proteins; the primary structural component of the PCM matrix is CDK5RAP2 in human cells, Centrosomin (Cnn) in *Drosophila*, and SPD-5 in *C. elegans* (Woodruff et al., 2014). During mitotic entry, the PCM matrix expands in a process referred to as centrosome maturation (Palazzo et al., 2000). Mitotic transition-coupled PCM expansion is under control of Polo-like kinase 1 (PLK1; (Cabral et al., 2019; Conduit et al., 2014; Dobbelaere et al., 2008; Haren et al., 2009; Lane and Nigg, 1996; Lee and Rhee, 2011; Woodruff et al., 2015)). In *Drosophila* and *C. elegans*, PLK1-stimulated PCM expansion has been shown to occur via isotropic incorporation of additional PCM matrix molecules (Conduit and Raff, 2015; Laos et al., 2015). In *Drosophila*, PCM matrix expansion occurs through a phospho-regulated self-interaction between Cnn molecules in which the CM2 (Centrosomin motif 2) motif at the C-terminus of one Cnn molecule interacts with an internal region in a second Cnn molecule that comprises a leucine zipper with adjacent PLK1 phosphorylation sites (Citron et al., 2018; Conduit et al., 2014; Feng et al., 2017). In *C. elegans*, a similarly positioned internal set of PLK1 sites has been identified that controls SPD-5 self-assembly (Woodruff et al., 2015).

The increase in microtubule nucleation capacity that accompanies centrosome maturation has been assumed to be a passive consequence of PCM matrix expansion. The PCM matrix docks nucleating complexes containing the specialized tubulin isoform γ-tubulin (Moritz et al., 1995; Moritz et al., 1998; Schnackenberg et al., 1998). The large ring-shaped γ-tubulin-containing complexes on the PCM matrix are formed via the lateral association of smaller Y-shaped hetero-tetrameric small complexes (γTuSCs); each γTuSC contains two molecules of γ-tubulin supported by a pair of structurally similar γ-tubulin-interacting proteins (e.g. GCP2 and GCP3 in humans) (Consolati et al., 2020; Kollman et al., 2011; Kollman et al., 2010; Liu et al., 2020; Oegema et al., 1999; Wieczorek et al., 2020b). The human and *Drosophila* PCM matrix proteins, CDK5RAP2 and Cnn, recruit γ-tubulin complexes via a conserved Centrosomin Motif 1 (CM1) in their N-termini (Choi et al., 2010; Fong et al., 2008; Samejima et al., 2008; Zhang and Megraw, 2007). Recent work has shown that a set of helices in the N-terminal domain of some γ-tubulin complex proteins intercalates with 3-alpha helix microproteins of the Mozart family, and that this intercalation can be important for interaction with tethering factors (Huang et al., 2020; Wieczorek et al., 2020a). For example, the CM1 region of CDK5RAP2 was shown to form a short parallel coiled-coil that interacts with a Mozart (MZT2) intercalated GCP2 subunit in the large γ-tubulin ring complex (Wieczorek et al., 2020a). Consistent with the idea that microtubule nucleating capacity increases passively as the PCM matrix expands, interaction of γ-tubulin complexes with the CM1 motif has not been proposed to be directly regulated.

Here, we use the *C. elegans* embryo to test the model that increased microtubule nucleation by mitotic centrosomes is a passive consequence of PLK1-triggered PCM expansion. Contrary to this idea, we find that PLK1 has two separable functions in centrosome maturation and controls the generation of γ-tubulin complex docking sites independently of PCM expansion. We show that simultaneously disrupting both processes by mutating critical PLK-1 target sites on the PCM matrix fully accounts for the actions of centrosomal PLK1 in transforming the centrosome during mitotic entry. Our results explain how PLK-1 activity remodels the centrosome and indicate that generation of phospho-regulated γ-tubulin docking sites on the PCM matrix is critical for spindle assembly and chromosome segregation.

## RESULTS

### PCM matrix expansion does not fully account for the function of centrosomal PLK-1

Across metazoans, PLK1 drives PCM matrix expansion during mitotic entry. In *C. elegans*, recruitment of PLK-1 to centrosomes requires an essential phospho-dependent docking site, thought to be generated by CDK-1, in the *C. elegans* CEP192 homolog, SPD-2 (**Fig. 1A**; (Decker et al., 2011)). Centrosomal PLK-1 promotes PCM expansion by phosphorylating residues, including S653 & S658, in the middle region of the PCM matrix component SPD-5 that are essential for its self-assembly (**Fig. 1A**; (Woodruff et al., 2015)). To test the idea that the essential function of centrosomal PLK-1 is to promote PCM matrix expansion, we compared the consequences of preventing the centrosomal targeting of PLK-1 to the consequences of blocking PCM matrix expansion. To disrupt PLK-1 targeting, we introduced single copy RNAi-resistant transgenes encoding wild-type (WT) or PLK-1 docking mutant (PD^mut^) SPD-2 (**Fig. S1A**) into embryos expressing *in situ*-tagged PLK-1::GFP (to monitor centrosomal PLK-1) and mCherry::SPD-5 (to monitor PCM matrix dynamics). Imaging embryos expressing transgenic WT SPD-2 (after endogenous SPD-2 depletion) confirmed prior work showing that PLK-1 remains concentrated in a central centriolar focus while the SPD-5-based PCM matrix expands around the centrioles during mitotic entry ((Cabral et al., 2019; Magescas et al., 2019; Mittasch et al., 2020); **Fig. 1B, Video S1**). In embryos expressing PD^mut^ SPD-2, little to no PLK-1 was detected at centrosomes, despite normal recruitment to kinetochores on chromosomes, and the SPD-5 matrix failed to expand as the embryos entered mitosis (**Fig. 1B, Video S1**). Thus, in agreement with prior work (Decker et al., 2011), docking of PLK-1 on SPD-2 is required for SPD-5 matrix expansion during mitotic entry.

**Figure 1.**
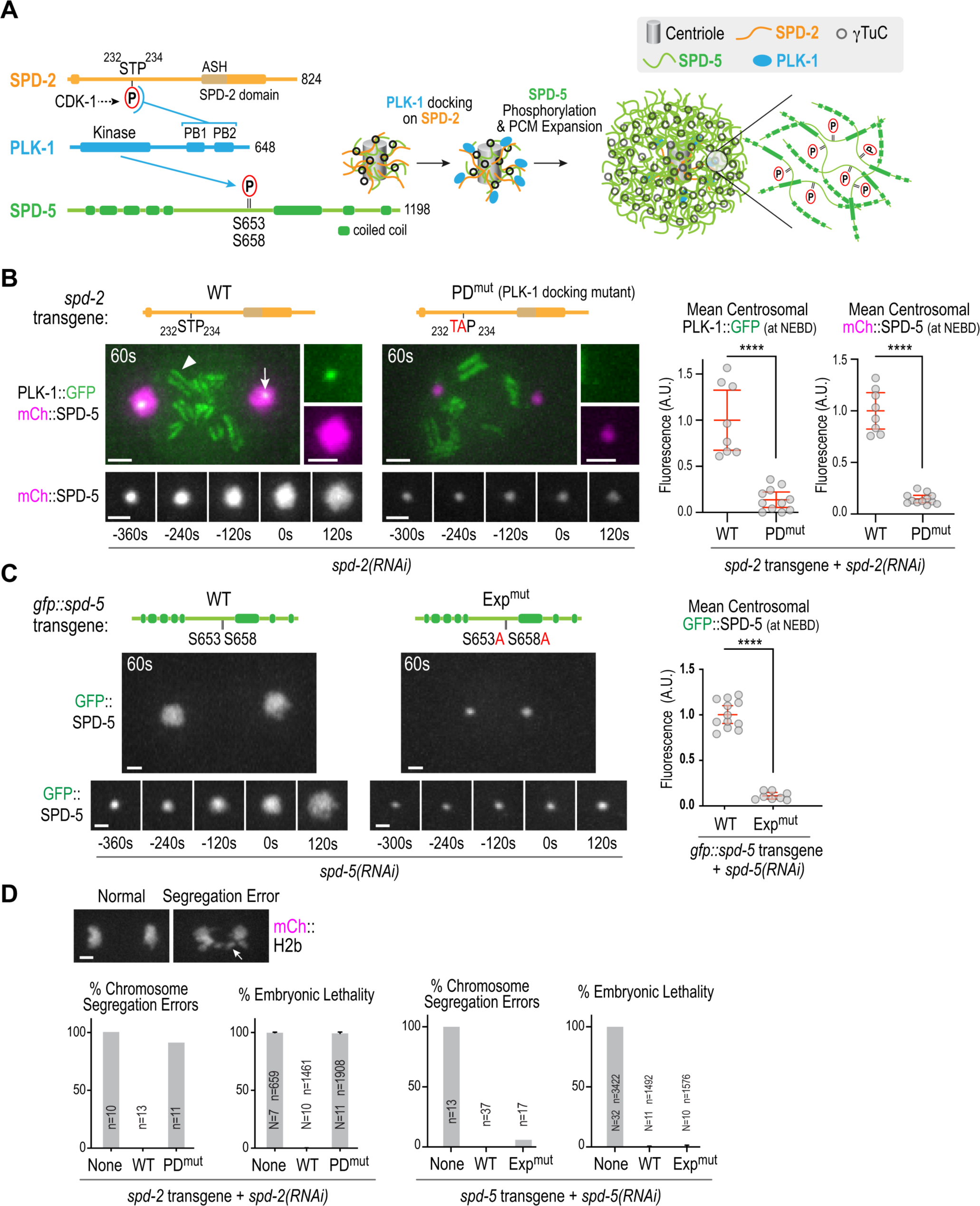
Pericentriolar matrix expansion does not fully account for the function of centrosome-localized PLK-1. (**A**) Current model for centrosome maturation *in C. elegans*. PLK-1 is recruited to centrosomes by binding of its C-terminal polo boxes (PB1 & PB2) to a site in SPD-2 (homolog of human CEP192) thought to be primed by Cdk1 (Thr233). PLK-1 controls PCM expansion by phosphorylating residues, including S653 and S658, in the central region of the PCM matrix component SPD-5 (analogous to CDK5RAP2 in mammals and Cnn in *Drosophila*) that enable its self-assembly. PCM expansion is assumed to passively increase the number of associated γ-tubulin complexes (*black rings*), and thus the ability of the centrosome to nucleate microtubules for spindle assembly. (**B, C**) Images of embryos expressing: (*B*) *in situ* GFP-tagged PLK-1 (*green*) and *in situ* mCherry-tagged SPD-5 (*magenta*) in the presence of wild-type (WT) or PLK-1 docking mutant (PD^mut^) SPD-2 after endogenous SPD-2 depletion; (*C*) wild-type (WT) or expansion-defective (Exp^mut^) GFP::SPD-5 after endogenous SPD-5 depletion. Graphs on right show quantification of centrosomal signals at NEBD. Error bars are mean with 95% CI. p-values are from t-tests. Times in seconds relative to NEBD are shown in top left of large panels and below the smaller panels, which show SPD-5 signal at one centrosome over time. In (*B*), PLK-1 localization to centrosomes (*arrow*) and kinetochores (*arrowhead*) in the presence of WT SPD-2, is indicated. (**D**) Analysis of chromosome segregation errors and embryonic lethality for the indicated conditions. (*top images*) Example of normal versus defective chromosome segregation, defined as visible chromatin bridges between anaphase chromosome masses in live imaging data. *n* refers to number of embryos imaged. For embryonic lethality analysis the mean ± SD is plotted, *N* refers to number of worms and *n* to number of embryos scored, respectively. All scale bars, 2 µm.

To compare the effect of preventing the centrosomal targeting of PLK-1 to the consequences of blocking SPD-5 matrix expansion by mutating PLK-1 target sites, we generated transgenes encoding GFP fusions with WT SPD-5 or SPD-5 in which two PLK-1 sites essential for expansion (S653, S658) are mutated to alanine (Exp^mut^ SPD-5; **Fig. S1B, Video S2**). Following endogenous SPD-5 depletion, Exp^mut^ SPD-5 exhibited an expansion defect comparable to that resulting from disrupting centrosomal PLK-1 targeting (**Fig. 1C**). However, despite their similar effects on SPD-5 matrix expansion, there was a striking difference in the phenotypes observed after preventing PLK-1 docking at centrosomes versus preventing SPD-5 matrix expansion (**Fig. 1D, Fig. S1C**). Penetrant chromosome segregation defects and embryonic lethality were observed in the former but not in the latter (**Fig. 1D**). These results indicate that the centrosomes in SPD-5 Exp^mut^ embryos, although small, support assembly of a spindle that segregates chromosomes; in addition, they suggest that PLK-1 controlled expansion of SPD-5 is not the only mechanism by which centrosomal PLK-1 acts during mitotic entry.

### A functional screen identifies putative PLK-1 target sites required to dock γ-tubulin complexes onto the PCM matrix

The results above suggested that there are other site(s) targeted by centrosomal PLK-1 that are important for spindle assembly and chromosome segregation. To identify such sites, we employed an unbiased strategy (Gomez-Cavazos et al., 2020), mutating putative PLK-1 target sites in SPD-5. In brief, we generated 7 strains expressing GFP::SPD-5 from single-copy RNAi-resistant transgenes in which regional clusters of candidate PLK-1 sites were mutated to alanines (**Fig. 2A, Fig. S1D,E**). The 7 cluster mutants, together with GFP-tagged WT and Exp^mut^ SPD-5, were analyzed following endogenous SPD-5 depletion (**Fig. 2A**). WT SPD-5 rescued the embryonic lethality resulting from endogenous SPD-5 depletion, as did Exp^mut^ SPD-5 and 6 of the alanine cluster mutants. Notably, the Cluster II mutant was associated with penetrant embryonic lethality. This result suggested that the three putative PLK-1 sites in Cluster II (S170, T178, T198) provide an essential function.

**Figure 2.**
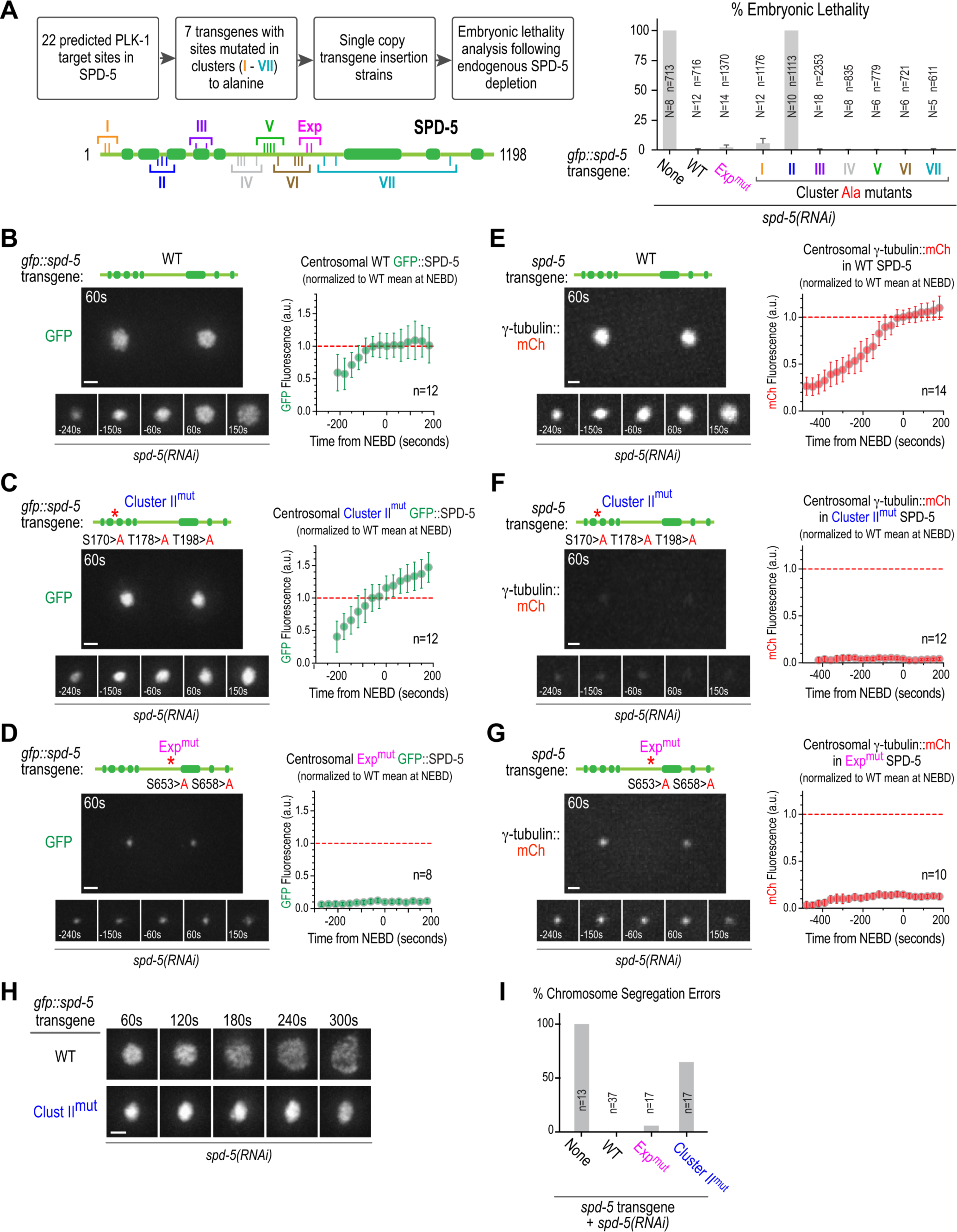
A putative PLK-1 target site cluster in SPD-5 required to dock γ-tubulin complexes onto the PCM matrix. (**A**) Flowchart of screening strategy. Predicted PLK-1 target sites were mutated in 7 clusters (*schematic*) and the consequences on embryonic lethality assessed. Graph on right plots embryonic lethality (mean ± SD) for the indicated conditions; *N* refers to number of worms and *n* to number of embryos scored. (**B**) – (**G**) Images and quantification of centrosomal intensity of GFP::SPD-5 variants (*B-D*) or of γ-tubulin::mCherry (*E-G*) in the presence of the indicated untagged SPD-5 variants; endogenous SPD-5 was depleted in all conditions. Centrosomal fluorescence was plotted after normalizing to the mean value at NEBD for WT GFP::SPD-5 (*B-D*) or for γ-tubulin::mCherry in the presence of WT SPD-5 (*E-G*). Error bars are the 95% CI. *n* refers to number of centrosomes imaged for each condition. (**H**) Individual centrosomes from the timelapse sequences of embryos expressing WT and Cluster II^mut^ GFP::SPD-5 shown in *B* and *C*. Timepoints highlight the PCM expansion that occurs during interval after anaphase onset when the WT GFP::SPD-5-based matrix disassembles under the influence of cortical pulling forces on matrix anchored microtubules. (**I**) Chromosome segregation analysis for the indicated conditions. *n* refers to the number of embryos imaged. The None, WT and Exp^mut^ data are reproduced from Figure 1D for comparison. Scale bars, 2 µm.

To assess the function of the putative PLK-1 target sites in Cluster II, we began by filming embryos expressing Cluster II mutant GFP::SPD-5. This analysis revealed that in contrast to Exp^mut^ SPD-5, Cluster II^mut^ SPD-5 did not block PCM expansion (**Fig. 2B-D, Video S2**). In fact, Cluster II^mut^ GFP::SPD-5 continued to accumulate at centrosomes even after levels of WT SPD-5 had plateaued. Despite the increased centrosomal signal, mitotic PCM assembled from Cluster II^mut^ GFP::SPD-5 remained more compact than PCM assembled from WT GFP::SPD-5; this was particularly evident after anaphase onset (**Fig. 2H, Video S2**) when the PCM matrix is subjected to forces that pull on PCM-anchored microtubules and promote PCM dispersal (Enos et al., 2018; Magescas et al., 2019; Mittasch et al., 2020). These results raised the possibility that PCM matrix assembled by Cluster II^mut^ SPD-5 is defective in the generation of PCM-anchored microtubules.

The PCM matrix nucleates and anchors microtubules in part due to bound γ-tubulin complexes. Quantitative monitoring of the centrosomal localization of mCherry tagged γ-tubulin (TBG-1 in *C. elegans*) revealed that, despite robust PCM matrix expansion, there was a dramatic reduction in the recruitment of γ-tubulin to the Cluster II^mut^ SPD-5 matrix (**Fig. 2E,F; Video S3**). Exp^mut^ SPD-5 also reduced γ-tubulin recruitment, but this was expected due to the reduction in PCM matrix size (**Fig. 2G, Video S3**). Consistent with normal PCM matrix assembly, neither PLK-1 nor SPD-2 centrosomal localization was affected in Cluster II^mut^ SPD-5 embryos (**Fig. S1F**). Finally, unlike Exp^mut^ SPD-5, Cluster II^mut^ SPD-5 was associated with significant defects in chromosome segregation (**Fig. 2I**). These segregation defects likely reflect problems in spindle assembly and underlie the embryonic lethality observed with Cluster II^mut^ SPD-5 (**Fig. 2A**).

Taken together, the above results suggest that PLK-1 independently controls expansion of the PCM matrix and the generation of γ-tubulin docking sites on the PCM matrix by phosphorylation of two different regions of SPD-5.

### PLK-1 phosphorylation promotes a direct interaction between SPD-5 and γ-tubulin complexes

The *in vivo* approaches described above identified a set of putative PLK-1 target sites in the SPD-5 N-terminus (S170, T178, T198) that are essential for γ-tubulin complex recruitment to the PCM matrix. To assess whether PLK-1 phosphorylation at these sites promotes a direct interaction between SPD-5 and γ-tubulin complexes, we established a biochemical reconstitution system. In *C. elegans*, γ-tubulin complexes (here termed Ce γTuCs) are composed of γ-tubulin and two γ-tubulin complex proteins (GCPs), GIP-1/GCP3 and GIP-2/GCP2 (**Fig. 3A**; (Hannak et al., 2001; Sallee et al., 2018)). *C. elegans* also has one homolog of the Mozart family of γ-tubulin complex associated microproteins (MZT-1), which is required to recruit γTuCs to the mitotic PCM (Sallee et al., 2018). This prior work, conducted in intestinal cells, also showed that although the γTuCs at interphase centrioles and the apical non-centrosomal microtubule organizing centers that form in this cell type normally contain MZT-1, MZT-1 is not required for γTuC targeting to these other locations (Sallee et al., 2018); thus, there are different γTuC targeting modalities, only some of which rely on MZT-1. Using a human cell expression system, we co-expressed the 3 core γTuC subunits (γ-tubulin, GIP-1, GIP-2) with and without MZT-1 (**Fig. 3A**) In the absence of GIP-2, GIP-1 and γ-tubulin were poorly expressed (**Fig. 3A**), indicating that association of the three core complex subunits is important for their stability. However, the three core subunits (γ-tubulin, GIP-1 and GIP-2) could be purified independently of MZT-1 (**Fig. 3A**) as might be expected based on the prior *in vivo* work (Sallee et al., 2018).

**Figure 3.**
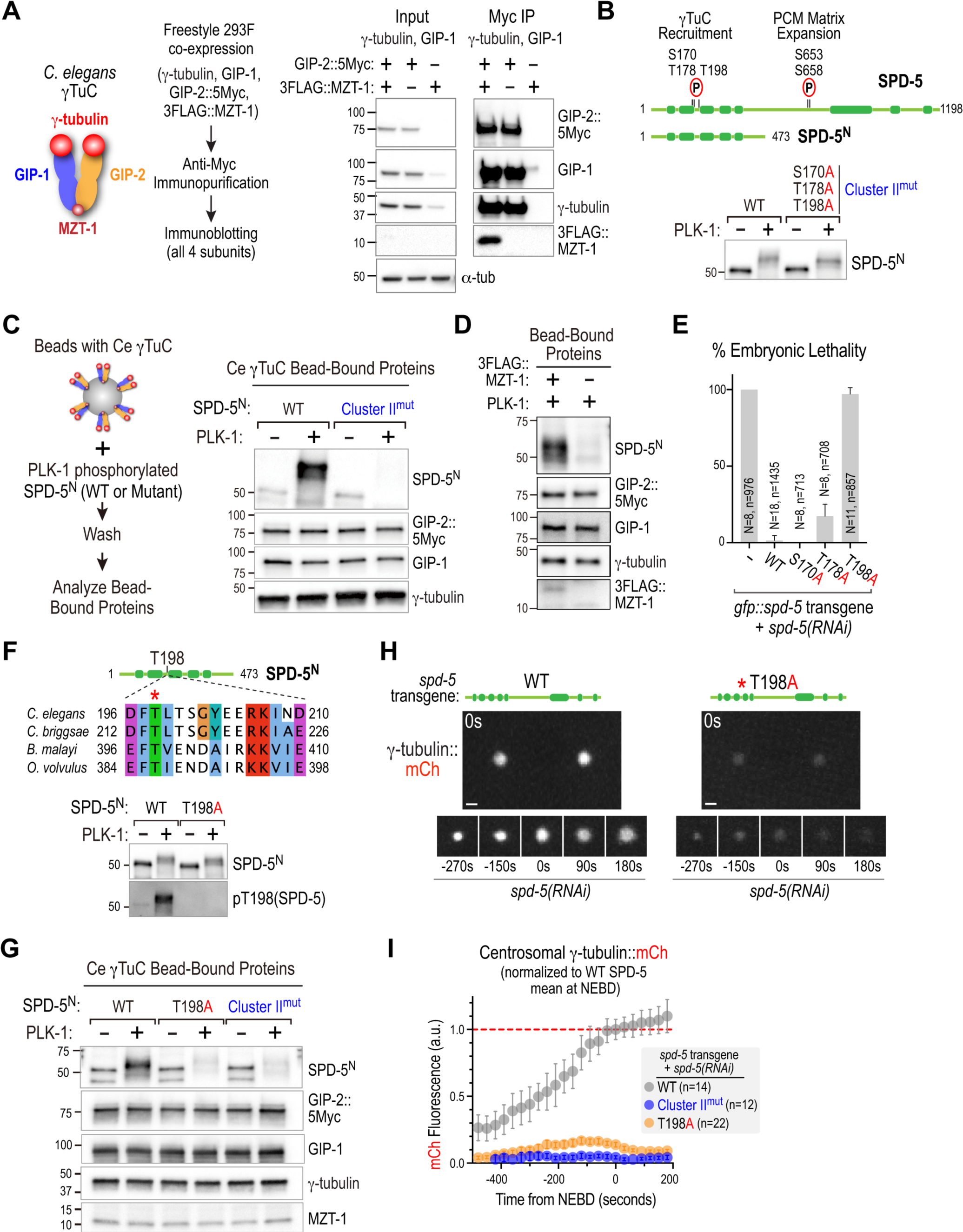
PLK-1 phosphorylation controls a direct interaction between SPD-5 and the γ-tubulin complex. (**A**) Reconstitution of the *C. elegans* γ-tubulin complex (*schematic*) by co-expression in mammalian cells (*flowchart in middle*). Plasmids encoding γ-tubulin, GIP-1 and GIP-2 (related to GCP3 and GCP2), and MZT-1 (related to Mozart) were co-transfected and the Myc tag on GIP-2 was used to immunopurify the complex. Immunoblots on right show Input (*left blot*) and anti-Myc immunopurified material (*right blot*) for the indicated conditions. (**B**) Schematic of PLK-1 sites in SPD-5 implicated in γ-tubulin recruitment (Cluster II; S170,T178,T198) and PCM matrix expansion (S653,S658). A SPD-5 N-terminal fragment (SPD-5^N^) that lacks the PCM expansion sites was bacterially expressed and purified. Blot shows effect of incubation of SPD-5^N^ variants with or without PLK-1 in presence of ATP for 1hr at 23°C. (**C**) (*left*) Schematic of analysis of interaction of γ-tubulin complexes, immobilized on beads, with SPD-5^N^ (either wildtype or Cluster II^mut^) pre-incubated with or without PLK-1. Blots of bead-bound components after incubation and isolation are shown on the right; FLAG-tagged MZT-1 was present but was not blotted here. (**D**) Analysis of interaction of PLK-1-phosphorylated SPD-5^N^ with γ-tubulin complexes reconstituted with or without FLAG-tagged MZT-1. (**E**) Graph of embryonic lethality (mean ± SD) for the indicated conditions, *N* refers to number of worms and *n* to number of embryos scored. (**F**) (*top*) Sequence alignment of divergent nematodes highlighting potential conservation of the critical T198 site. (*bottom*) Indicated SPD-5^N^ variants incubated with or without PLK-1 and immunoblotted to detect total SPD-5 (*top*) and phosphorylation of T198 (*bottom*), using a phospho-specific antibody (pT198). (**G**) Analysis of γ-tubulin complex binding, conducted as in (*C*), for the indicated conditions. (**H**)-(**I**) Representative images and quantification of γ-tubulin::mCherry recruitment to centrosomes for the indicated conditions. Error bars are the 95% CI. Scale bar, 2 µm. Data plotted for WT and Cluster II^mut^ SPD-5 is reproduced from Figures 2E & F for comparison.

In parallel to reconstituting Ce γTuC, we expressed and purified an N-terminal fragment of SPD-5 (amino acids 1-473, SPD-5^N^; **Fig. S2A**); the fragment was designed to lack the region of SPD-5 whose phosphorylation controls PCM matrix expansion (**Fig. 3B**). To assess if Cluster II sites are phosphorylated by PLK-1, we incubated the recombinant SPD-5 fragment with constitutively-active PLK-1 (**Fig. 3B**). SPD-5^N^ mobility was substantially retarded following incubation with PLK-1, with the extent of retardation being slightly reduced by mutation of the Cluster II residues (**Fig. 3B**). This result suggested that Cluster II sites are targets of PLK-1 kinase activity *in vitro*, with additional sites in SPD-5^N^ also being targeted by PLK-1.

To determine if PLK-1 phosphorylation of Cluster II sites controls the ability of the SPD-5 N-terminus to interact with γ-tubulin complexes, we immobilized reconstituted γTuCs on beads, mixed the beads with unphosphorylated or PLK-1 phosphorylated recombinant SPD-5^N^, and analyzed bead-bound proteins (**Fig. 3C**). PLK-1 phosphorylation substantially enhanced the association of SPD-5^N^ with γTuC-coated beads. Notably, the ability of PLK-1 to enhance interaction of SPD-5^N^ with the γTuC was abrogated by mutation of the Cluster II residues (**Fig. 3C**). The ability of the γTuC to interact with phosphorylated SPD-5^N^ also required MZT-1 (**Fig. 3D**), consistent with the prior work showing that MZT-1 is required for γTuCs to be recruited to the mitotic PCM in vivo (Sallee et al., 2018). We conclude that MZT-1 is an integral γTuC component required for its PLK-1-dependent docking on the SPD-5 PCM matrix. Consistent with this, the Cluster II mutant had the same effect on the centrosomal accumulation of *in situ*-tagged GFP::MZT-1 as it did on γ-tubulin::mCherry (**Fig. S2B,C; Fig. 2F**). Thus, γ-tubulin complexes with integrated MZT-1 are recruited to SPD-5 in the PCM that is phosphorylated on Cluster II by centrosomal PLK-1.

### A single PLK-1 target site controls the majority of γ-tubulin complex recruitment to the mitotic PCM matrix

Sequence alignments of SPD-5s from related nematode species indicated that within Cluster II T178 and T198 are conserved, whereas S170 is not (*not shown*). To evaluate the importance of these residues, we analyzed embryonic lethality following their individual mutation. Mutation of S170 had no effect, mutation of T178 resulted in mild (∼17%) lethality and mutation of T198 resulted in penetrant (>95%) lethality (**Fig. 3E**). Consistent with this penetrant lethality, sequence analysis of highly divergent nematode species (∼400 Mya apart) suggested T198 is conserved (**Fig. 3F**). A phospho-specific antibody raised against pT198 site confirmed phosphorylation at this site by PLK-1 *in vitro* (**Fig. 3F**) and mutation of T198 alone significantly compromised the ability of PLK-1 to promote association of SPD-5^N^ with γTuCs *in vitro* (**Fig. 3G**) and the ability of the SPD-5 matrix to recruit γTuCs *in vivo* (**Fig. 3H,I; Fig. S2B,C; Video S3**). These data highlight T198 as being a critical PLK-1 target site whose phosphorylation controls γ-tubulin complex docking on the mitotic PCM.

We consistently observed slightly more γ-tubulin complex recruitment in embryos expressing T198A relative to Cluster II^mut^ SPD-5 (**Fig. 3H,I; Fig. S2B,C; Video S3**), suggesting a minor contribution to γ-tubulin complex recruitment by one of the other sites in Cluster II, most likely T178 based on conservation and lethality measurements. In contrast to Cluster II^mut^ SPD-5, which failed to support embryonic viability when untagged or GFP-tagged, the residual γ-tubulin complex recruitment supported by T198A SPD-5 was sufficient for embryonic viability when untagged but not when GFP-tagged (**Fig. S2D**). Overall, these results suggest that phosphorylation by PLK-1 of SPD-5 T198 plays a major role in γ-tubulin complex docking on the PCM, with phosphorylation on T178 likely playing a minor role. These results also reinforce the fact that relatively small amounts of centrosomal γ-tubulin (<20% of normal levels) are sufficient to support embryonic viability.

### PLK-1 controlled γ-tubulin complex docking and PCM expansion collaboratively support spindle assembly

The analysis above indicates that PLK-1 has two separable functions during centrosome maturation; it controls γ-tubulin complex docking by phosphorylating a region in the SPD-5 N-terminus and it promotes expansion of the PCM matrix by phosphorylating a region in the middle of SPD-5. To address to what extent these two functions, both working through a common substrate, account for the action of centrosomal PLK-1, we analyzed spindle assembly and chromosome segregation. To define the action of centrosomal PLK-1 in spindle assembly, we analyzed PD^mut^ SPD-2 in a strain with labeled microtubules and chromosomes (**Fig. 1B**). Quantitative analysis of centrosome separation and centrosomal β-tubulin intensity revealed that disrupting centrosomal PLK-1 recruitment significantly compromised microtubule assembly around centrosomes, centrosome separation, and chromosome segregation (**Fig. 4A,C; Video S4**).

**Figure 4.**
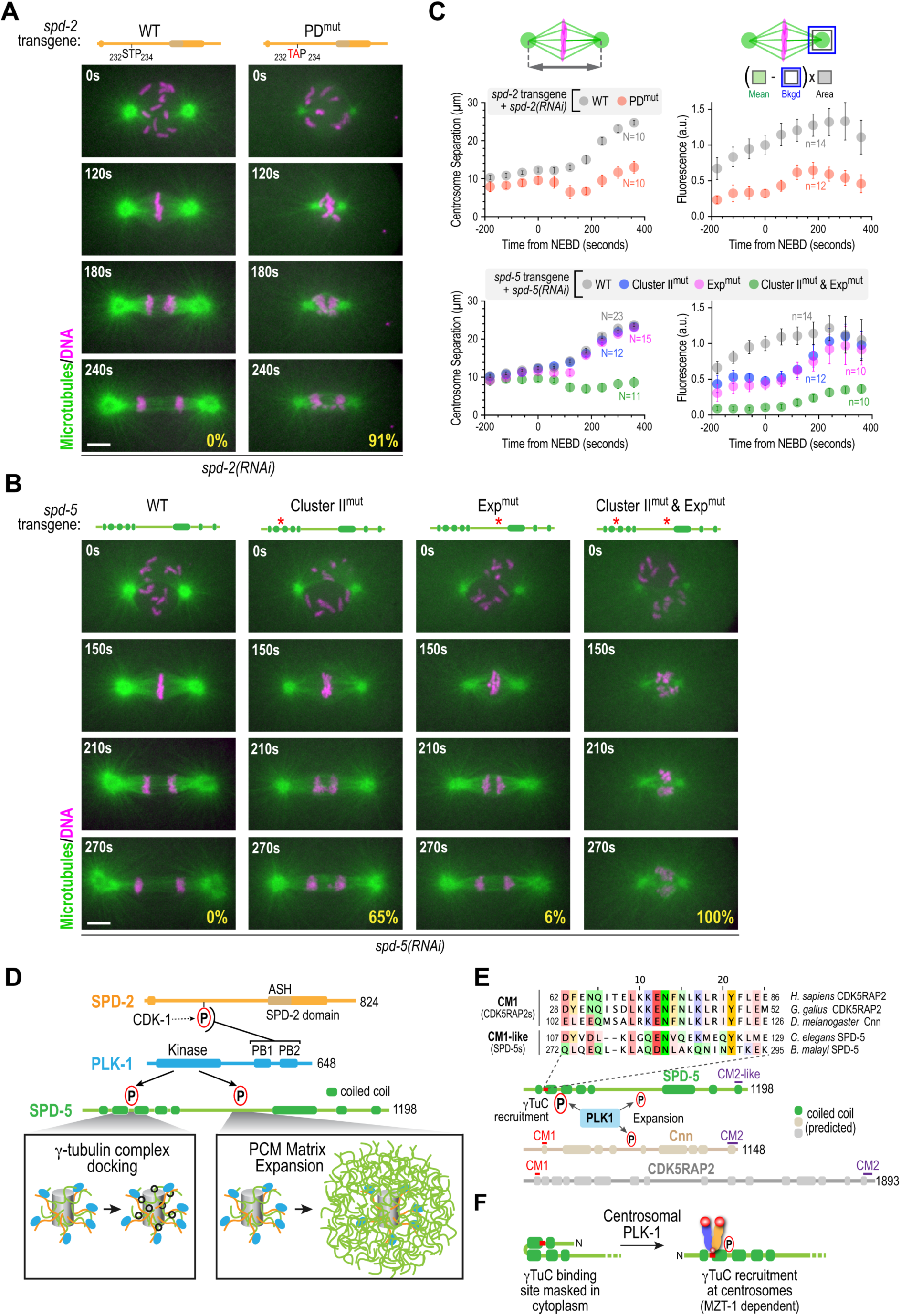
Independent phospho-regulation of SPD-5-anchored γ-tubulin recruitment and PCM expansion fully accounts for the function of PLK-1 in centrosome maturation. (**A,B**) Images of GFP::β-tubulin (*green*) and mCherry::histone (*magenta*) in embryos expressing the indicated untagged SPD-2 (*A*) or SPD-5 (*B*) variants with endogenous SPD-2 (*A*) or SPD-5 (*B*) depleted. Times in the top left of each panel are seconds after NEBD. Percentages of chromosome segregation errors are in the bottom right of each panel. Chromosome segregation data for conditions other than the double mutant is the same as that shown in Figures 1D and 2I and is shown here for comparison. Scale bars are 5 µm. (**C**) Quantification of centrosome separation (*left*) or centrosomal β-tubulin intensity (*right*) for the indicated conditions. Error bars are the 95% confidence interval. *N* refers to number of embryos and *n* to number of centrosomes scored. (**D**) Schematic depicting independent phospho-regulation by centrosome-localized PLK-1 of γ-tubulin recruitment and PCM expansion through phosphorylation of distinct regions of SPD-5. (**E**) Sequence alignments supporting nematode SPD-5s being divergent orthologs of CDK5RAP2 and Cnn. SPD-5s from distantly related nematodes (∼400 Mya apart) have a CM1-like motif N-terminal to the key phospho-regulated sites identified here (see **Fig. 3F**). SPD-5 also has a CM2-like motif at its C-terminus (**Fig. S3C**) and a central PReM-like region that contains the PLK-1 sites previously shown to control its assembly (**Fig. S3D**). (**F**) Speculative model for mechanism of phospho-regulated γ-tubulin complex recruitment. The CM1-like recruitment site is masked in unphosphorylated cytoplasmic SPD-5. Phosphorylation at centrosomes unmasks the binding site, leading to γ-tubulin complex recruitment specifically on pericentriolar matrix-assembled SPD-5.

When γ-tubulin complex docking or PCM matrix expansion were disrupted individually using the SPD-5 Cluster II or Exp mutants, the phenotypes were significantly milder than disruption of centrosomal PLK-1. In both cases, centrosomal microtubule intensity was reduced, albeit to a lesser degree than when centrosomal PLK-1 recruitment was disrupted; however, centrosome separation kinetics were not significantly altered (**Fig. 4B,C; Video S4**). To assess if simultaneously disrupting PLK-1 driven γ-tubulin complex docking and PCM expansion recapitulated the effect of preventing PLK-1 centrosomal recruitment, we generated a double Cluster II^mut^ & Exp^mut^ SPD-5. The double mutant SPD-5 exhibited similar severity defects—in centrosomal microtubule intensity, centrosome separation and chromosome segregation—as PD^mut^ SPD-2 (**Fig. 4B,C; Video S4**). Thus, the severe spindle assembly defect resulting from disrupting the centrosomal targeting of PLK-1 can be reconstituted by preventing the phosphorylation of two sets of PLK-1 target sites in the substrate SPD-5 that independently control γ-tubulin complex docking and PCM matrix expansion.

Despite essentially identical centrosome separation kinetics, an interesting difference between the individual Cluster II^mut^ and Exp^mut^ phenotypes was the extent of chromosome missegregation; chromosome segregation defects were significantly higher in the presence of Cluster II^mut^ versus Exp^mut^ SPD-5 (**Fig. 4B**, see also **Fig. 2I**). We suspect that this difference may arise from the nature of the spindle microtubules in these two conditions. In SPD-5 Exp^mut^ embryos, spindle microtubules are likely generated by SPD-5 anchored γ-tubulin complexes as in control embryos, but there are fewer microtubules because of the lack of PCM expansion. The origin of the microtubules in SPD-5 Cluster II^mut^ embryos, in which the PCM matrix is severely compromised in its ability to dock γ-tubulin complexes, is less clear. One possibility is that the remaining microtubules are generated by γ-tubulin complexes recruited from the cytoplasm that cap microtubule minus ends but are not PCM anchored. As PCM assembly is unaffected in Cluster II^mut^ embryos (**Fig. 2C**), a second possibility is that the remaining spindle microtubules are nucleated by γ-tubulin-independent mechanisms involving other PCM-localized factors and/or the ability of centrosomes to locally concentrate soluble tubulin (Baumgart et al., 2019; Woodruff et al., 2017). A serendipitous observation leads us to favor the first possibility. When assessing the effect of Cluster II^mut^ SPD-5 on γ-tubulin complex recruitment, we analyzed the recruitment of *in situ* GFP-tagged GIP-2 which, like γ-tubulin::mCh and GFP::MZT-1, failed to localize to the mitotic PCM in Cluster II^mut^ SPD-5 embryos (**Fig. S3A**). However, there was a strong synthetic phenotype observed when combining Cluster II^mut^ SPD-5 with in situ GFP-tagged GIP-2; specifically, centrosome separation was inhibited in this strain to a degree even more severe than when centrosomal PLK-1 recruitment was disrupted (**Fig. S3B**). This genetic interaction strongly suggests that microtubule and spindle assembly in SPD-5 Cluster II^mut^ embryos may depend on microtubules nucleated by non-PCM anchored γ-tubulin complexes rather than on microtubules nucleated by γ-tubulin-independent mechanisms.

## DISCUSSION

Here we addressed the mechanism by which PLK1 transforms the centrosome during mitotic entry to support spindle assembly. We tested the model that the mitotic expansion of the PCM matrix, under control of centrosomally-targeted PLK1, passively increases microtubule nucleation capacity at centrosomes. Our results suggest that PCM expansion is not sufficient to account for the function of centrosomal PLK1 because PLK1 independently controls γ-tubulin complex docking onto the PCM matrix (**Fig. 4D**). Thus, centrosome maturation during mitotic entry is accomplished via the action of two separable PLK1-dependent processes, PCM matrix expansion and γ-tubulin complex docking, which collaborate to increase the nucleating capacity of centrosomes for spindle assembly.

### PCM matrix components: conserved modes of regulation?

*C. elegans* SPD-5 shares many properties with the PCM matrix components Cnn and CDK5RAP2. Expansion of the Cnn-based PCM matrix in *Drosophila* occurs through PLK1 phospho-regulated self-assembly (Citron et al., 2018; Conduit et al., 2014; Feng et al., 2017). Sequence analysis indicated that nematode SPD-5s have regions with homology to the domains of Cnn that are critical for this PLK1 regulated self-assembly (**Fig. S3C,D**). A hallmark feature of *Drosophila* Cnn and human CDK5RAP2 is the presence of a conserved CM1 motif, which directly interacts with γ-tubulin-complexes (Choi et al., 2010; Fong et al., 2008; Samejima et al., 2008; Zhang and Megraw, 2007). Structural work has shown that the CM1 region of human CDK5RAP2 forms a short parallel coiled-coil that interacts with a MZT2-intercalated GCP2 subunit in the γ-tubulin complex (Wieczorek et al., 2020a). The CM1 motif interaction with γ-tubulin complexes has not been found to require phosphorylation. Sequence analysis suggests presence of a divergent CM1-like domain in nematode SPD-5s, ∼70 amino acids N-terminal to the critical PLK-1 target sites identified here (**Fig. 4E**). Based on structural work with human proteins and given that the interaction we characterize between PLK-1 phosphorylated SPD-5 and γ-tubulin complexes depends on *C. elegans* Mozart (MZT-1), we speculate that PLK-1 phosphorylation generates γ-tubulin docking sites by unmasking the CM1-like domain on SPD-5 molecules that are incorporated into the PCM matrix (**Fig. 4F**). In this model, PLK-1 phosphorylation releases autoinhibition intrinsic to the SPD-5 N-terminus to expose the CM1-like γ-tubulin docking site on SPD-5. An appealing feature of this model is that it would constrain association with γ-tubulin complexes to SPD-5 assembled into the PCM matrix, in the vicinity of concentrated PLK-1 activity, instead of globally in the cytoplasm.

We note that in human cells phosphorylation by LRRK1 of Ser140 of CDK5RAP2, 54 amino acids distal to the CM1 domain, has been proposed to regulate interaction with γ-tubulin complexes (Hanafusa et al., 2015). The identified LRRK1 target site (S140) is not conserved even in vertebrates; however, there are PLK1 sites in the region following the core CM1 motif in vertebrate CDK5RAP2 and *Drosophila* Cnn that may act in a manner similar to the sites we describe here for SPD-5. Future experimental analysis in different organisms, together with continuing analysis of SPD-5 in *C. elegans* based on the sequence comparisons shown here, will be important to test these ideas.

Overall, the results presented here explain how centrosome-targeted PLK-1 remodels the centrosome for spindle assembly and reveal tight phospho-regulation of γ-tubulin docking on the PCM matrix during the transition from interphase to mitosis. We suggest that similar phosphoregulation may be employed in different contexts by diverse γ-tubulin complex tethering factors to ensure the spatial regulation of microtubule nucleation.

## ACKNOWLEDGMENTS

We thank J. Woodruff for purified PLK-1, J. Feldman for the GFP::MZT-1 strain, members of the Oegema and Desai labs for helpful discussions, and Franz Meitinger for critical reading. This work was supported by an NIH grant to K.O. (GM074207). M.O. was supported by the Japan Society for the Promotion of Science. J.S.G-C was supported by the University of California, San Diego Cancer Cell Biology Training Program (T32 CA067754) and Ruth L. Kirschstein Postdoctoral Individual National Research Service Award (F32GM125347). A.D. and K.O. acknowledge salary support from the Ludwig Institute for Cancer Research.

## AUTHOR CONTRIBUTIONS

Conceptualization, K.O., A.D., M.O.; Methodology, K.O, A.D., M.O.; Resources, D.W., S.W., J.L.H., J. S. G-C; Investigation, M.O., Z.Z.; Writing – Original Draft, K.O, A.D., M.O.; Writing – Review & Editing, K.O., A.D., M.O.; Funding Acquisition, K.O., A.D., M.O.; Visualization, K.O., A.D., M.O.; Supervision, K.O., A.D.

## FIGURES, FIGURE SUPPLEMENTS, AND LEGENDS

**Figure S1 (Related to Figures 1 and 2).**
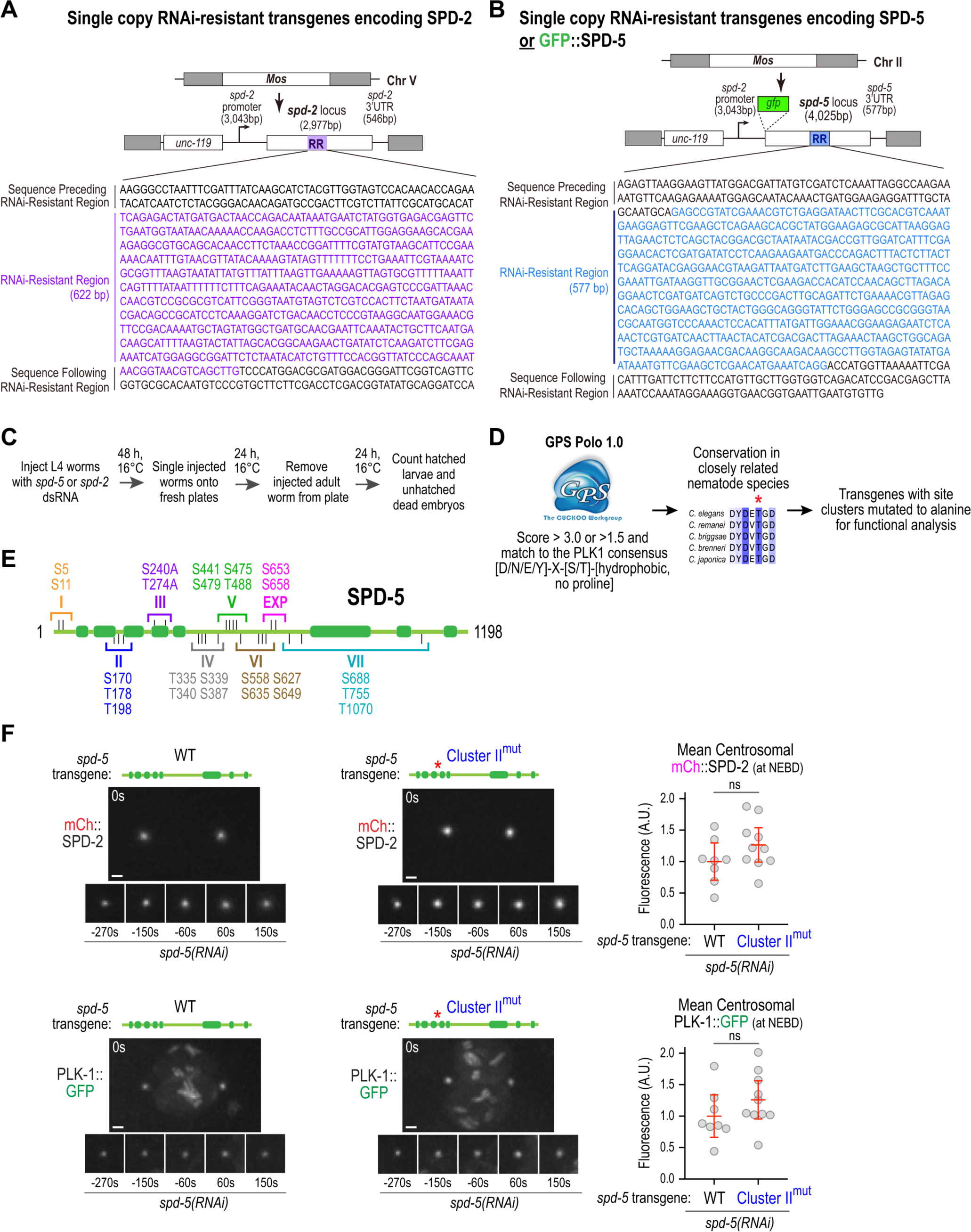
Transgene replacement systems for SPD-2 and SPD-5, method for identifying putative PLK-1 sites in SPD-5, and analysis of PLK-1 and SPD-2 centrosomal localization in Cluster II mutant SPD-5. (**A**) Single-copy RNAi-resistant transgene insertion system used to express SPD-2 variants. Purple region marked “RR” was altered in nucleotide but not in protein coding sequence to make the transgene-encoded product resistant to dsRNAs targeting the equivalent region of endogenous *spd-2*. (**B**) Single-copy RNAi-resistant transgene insertion system used to express untagged and GFP-tagged SPD-5 variants. Blue region marked “RR” was altered in nucleotide but not coding sequence to make the transgene-encoded product resistant to dsRNAs targeting the equivalent region of endogenous *spd-5*. Note that due to repetitive architecture of its promoter region, the *spd-2* promoter was used to control expression of *spd-5* transgenes. (**C**) Protocol employed to assess embryonic lethality. L4 worms were injected with dsRNA targeting *spd-5* or *spd-2* and embryos laid 48-72h (at 16°C) after injection were scored for lethality. (**D**) Outline of procedure used to identify putative PLK-1 target sites in SPD-5. (**E**) Schematic depicting identified sites and regional clusters employed for mutagenesis. 7 transgenes harboring mutant clusters I-VII were used to generate single copy integration strains that were used for the embryonic lethality analysis shown in *Fig. 2A*. (**F**) Images of *in-situ* tagged mCherry::SPD-2 *(top)* and PLK-1::GFP *(bottom)* in embryos expressing WT or Cluster II^mut^ SPD-5 after endogenous SPD-5 depletion. Times in the top left of large panel and below smaller panels are in seconds relative to NEBD. Scale bars, 2 µm. Graphs on right show quantification of centrosomal mCherry::SPD-2 or PLK::GFP signal at NEBD. Error bars are mean with 95% CI. p-values are from t-tests.

**Figure S2 (Related to Figure 3).**
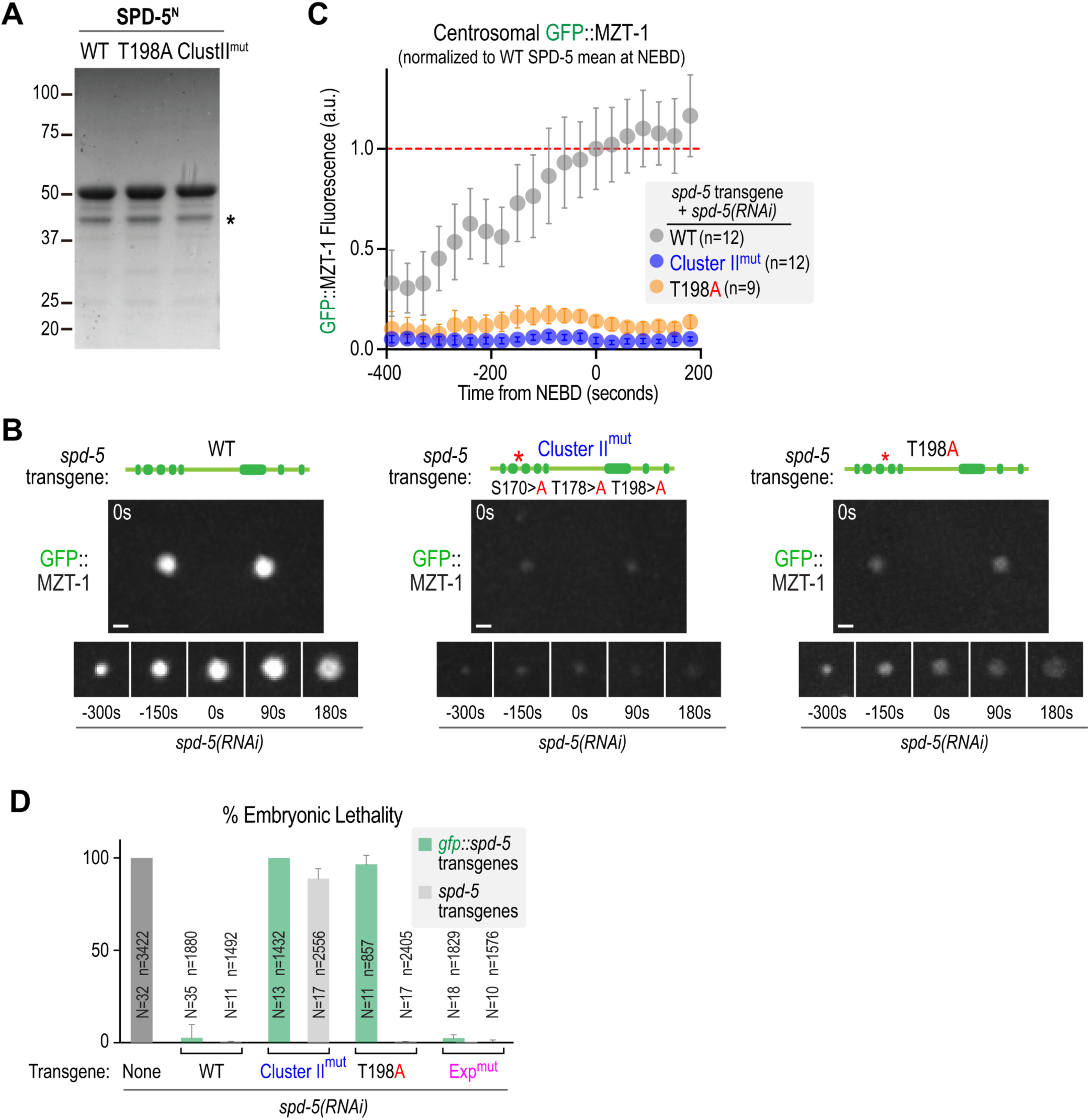
Analysis of GFP::MZT-1 localization and comparison of T198A SPD-5 transgenes with and without a GFP tag. (**A**) Coomassie blue stained gel showing the purified SPD-5 N-terminal fragments (amino acids 1-473, SPD-5^N^) used in the interaction assays. Asterisk marks a SPD-5^N^ breakdown product. (**B**) Images of *in-situ* tagged GFP::MZT-1 for the indicated conditions. Times shown in top left of large panel and below smaller panels are in seconds relative to NEBD. Scale bar, 2 µm. (**C**) Quantification of centrosomal GFP::MZT-1 signal. Values were normalized relative to the mean at NEBD in the presence of WT SPD-5. Error bars are the 95% CI. *n* refers to the number of centrosomes. (**D**) Embryonic lethality is plotted for the indicated conditions. The data for the non-gfp containing transgenes (None, WT and Exp^mut^) conditions is the same as that shown in Figure 1D and is reproduced here for comparison. *N* refers to the number of worms and *n* to the number of embryos scored. T198A mutant SPD-5 shows penetrant lethality when fused to GFP but not when untagged. No enhancement of lethality is observed when comparing GFP-fused and untagged Expansion-mutant SPD-5.

**Figure S3 (Related to Figure 4).**
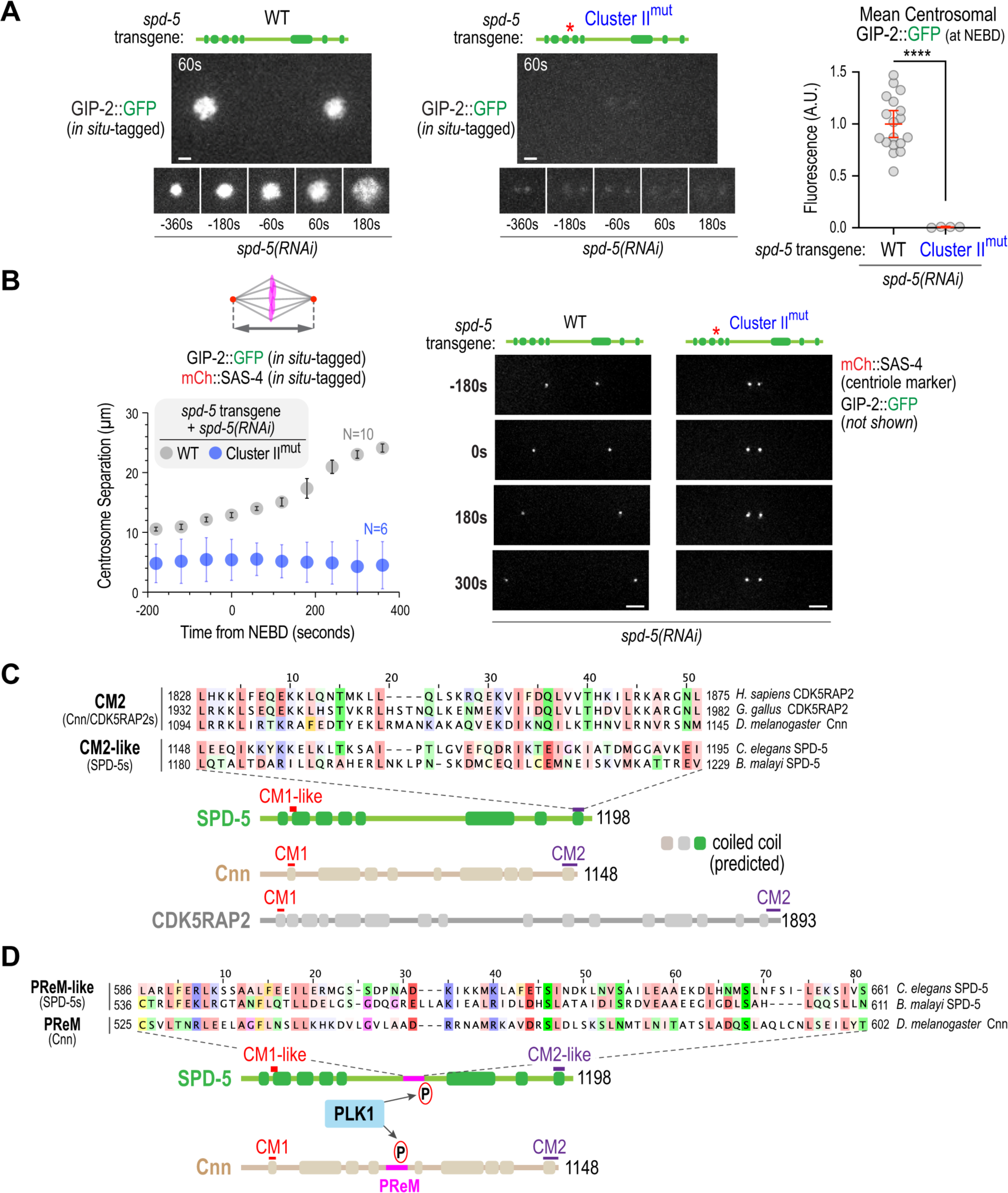
*In situ*-tagged GIP-2::GFP exhibits a synthetic spindle assembly phenotype when combined with Cluster II-mutant SPD-5. (**A**) Example images of GIP-2::GFP in the presence of WT or Cluster II-mutant SPD-5. Times in the top left of large panel and below smaller panels are seconds relative to NEBD. Scale bar, 2 µm. Error bars are mean with 95% CI. p-values are from t-tests. (**B**) Quantification of centrosome separation for the indicated conditions. Spindle length was measured using mCherry::SAS-4 that was also expressed in the analyzed strains. Times are seconds after NEBD. Error bars are the 95% CI. *N* refers to number of embryos imaged. (**C,D**) Nematode SPD-5s have regions with homology to the domains of *Drosophila* Cnn that mediates its PLK1-regulated self-assembly. Expansion of the Cnn-based PCM matrix occurs through a phospho-regulated self-interaction in which a conserved CM2 motif at the Cnn C-terminus interacts with an internal region of Cnn (called the phosphoregulated-multimerization or “PReM” domain) composed of a leucine zipper followed by a control region that enables assembly when phosphorylated by PLK1 (Citron et al., 2018; Conduit et al., 2014; Feng et al., 2017). (C) Sequence alignment suggesting the presence of a CM2-like domain at the SPD-5 C-terminus. (D) Sequence alignment showing the region of SPD-5 containing the PLK1 sites previously shown to control SPD-5 self-assembly (Woodruff et al., 2015). These sites follow a short a short leucine-rich region similar to that in the PReM region of *Drosophila* Cnn (Conduit et al., 2014).

## SUPPLEMENTAL VIDEO LEGENDS

**Video S1 (Related to Figure 1). Centrosomal PLK-1 is required for PCM expansion during mitotic entry**. Spinning disk confocal optics were used to image one-cell stage *C. elegans* embryos expressing *in situ*-tagged PLK-1::GFP (*green*) and mCherry::SPD-5 (*magenta*) in the presence of transgene-encoded wild-type (WT) or PLK-1 docking mutant (PD^mut^) SPD-2; endogenous SPD-2 was depleted. Times shown in bottom left are seconds after NEBD. 11 × 1 μm z-stacks were acquired every 60 seconds and maximal intensity projections were generated for each time frame. Playback rate is 2 frames per second. Embryos were filmed at 20°C. Scale bars, 10 µm. Related to Fig. 1B.

**Video S2 (Related to Figures 1 and 2). Putative PLK-1 target site Cluster II mutant SPD-5 did not block PCM expansion**. Spinning disk confocal optics were used to image one-cell stage *C. elegans* embryos expressing transgene-encoded GFP fusion with wild-type (WT), Cluster II mutant (Cluster II^mut^) or expansion mutant (Exp^mut^) SPD-5; endogenous SPD-5 was depleted. Times shown in bottom left are seconds after NEBD. 6 × 2 μm z-stacks were acquired every 30 seconds and maximal intensity projections were generated for each time frame. Playback rate is 2 frames per second. Embryos were filmed at 20°C. Scale bars, 10 µm. Related to Fig. 1C and 2B-D.

**Video S3 (Related to Figures 2 and 3). PLK-1 target site Cluster II in SPD-5 is required to dock γ-tubulin complexes onto the PCM matrix**. Spinning disk confocal optics were used to image one-cell stage *C. elegans* embryos expressing transgene encoded γ-tubulin::mCherry in the presence of transgene-encoded untagged wild-type (WT), Cluster II mutant (Cluster II^mut^), expansion mutant (Exp^mut^) or T198A SPD-5; endogenous SPD-5 was depleted. Times shown in bottom left are seconds after NEBD. 13 × 1 μm z-stacks were acquired every 30 seconds and maximal intensity projections were generated for each time frame. Playback rate is 2 frames per second. Embryos were filmed at 20°C. Scale bars, 10 µm. Related to Fig. 2E-G and 3H.

**Video S4 (Related to Figure 4). PLK-1-controlled γ-tubulin recruitment and expansion of SPD-5 account for the function of centrosome-localized PLK-1**. Spinning disk confocal optics were used to image one-cell stage *C. elegans* embryos expressing transgene-encoded GFP:: β-tubulin (*green*) and mCherry::histone (*magenta*) in the presence of transgene-encoded untagged wild-type (WT) or PLK-1 docking mutant (PD^mut^) SPD-2, or wild-type (WT), Cluster II mutant (Cluster II^mut^), expansion mutant (Exp^mut^) or Cluster II^mut^ & Exp^mut^ SPD-5; endogenous SPD-2 or SPD-5 was depleted. Times shown in bottom left are seconds after NEBD. 11 × 1µm z-stacks were acquired every 60 seconds for SPD-2 variants; 9 × 1 μm z-stacks were acquired every 30 seconds for SPD-5 variants. For microtubule imaging, the three z-planes containing the spindle poles were projected for each timepoint. For mCherry::histone, all z-planes were projected for each timepoint. Playback rate is 2 frames per second. Embryos were filmed at 20°C. Scale bars, 10 µm. Related to Fig. 4A, B.

## MATERIALS AND METHODS

### *C. elegans* strains

*C. elegans* strains (listed in Table S1) were maintained at 16°C. Single-copy transgenes were generated by using the transposon-based MosSCI method (Frokjaer-Jensen et al., 2008) to recombine them into specific chromosomal sites. Transgenes were cloned into pCFJ151 and injected into strains with specific Mos transposon insertions to recombine them into the ttTi5605 site on Chr II or the Uni V oxTi365 site on Chr V. Transgenes were generated by injecting a mixture of the pCFJ151-derived repairing plasmid containing the Cb-unc-119 selection marker and appropriate homology arms (50-100 ng/μL), transposase plasmid (pCFJ601 encoding the Mos1 transposase under the Peft-3 promoter, 50 ng/μL) and four plasmids encoding markers for negative selection against chromosomal arrays (pMA122 [Phsp-16.41::peel-1, 10 ng/μL], pCFJ90 [Pmyo-2::mCherry, 2.5 ng/μL], pCFJ104 [Pmyo-3::mCherry, 5 ng/μL] and pGH8 [Prab-3::mCherry, 10 ng/μL]) into strains EG6429 (ttTi5605, Chr II) or EG8082 (oxTi365, Chr V). After one week, the progeny of injected worms were heat-shocked at 34°C for 3 hours to induce the expression of PEEL-1 to kill worms containing extra chromosomal arrays. Moving worms without fluorescent markers were identified as candidates, and PCR across the junctions on both sides of the integration site was used to confirm transgene integration in their progeny.

### Generation of single-copy insertion transgenes

We had previously generated RNAi-resistant transgenes encoding SPD-2 (Shimanovskaya et al., 2014) and SPD-5 (Woodruff et al., 2015). The same replacement strategies (**Fig. S1A,B**) were used to generate the SPD-2 and SPD-5 encoding transgenes described here. Transgenes were generated by PCR amplification of their respective genomic loci. The *spd-2* transgene included 3043 bp region upstream of the start codon and 546 bp downstream of the stop codon. As the high level of repetitive sequence in the endogenous *spd-5* promoter made the single-copy insertion procedure inefficient, the *spd-5* transgenes are driven by the *spd-2* promoter and included 577 bp downstream of the stop codon. Segments of the *spd-2* and *spd-5* transgenes were modified as indicated (**Fig. S1A,B**) to make the transgenes RNAi-resistant without altering coding information. To generate the strains expressing transgenic GFP::SPD-5 with the individual S170A, T178A and T198A mutations (used in Fig. 3E and S2D), CRISPR was used as previously described (Hattersley et al., 2018) to directly introduce the mutations into the re-encoded region of the WT SPD-5 transgene. Worms from the strain OD2435 were injected with purified Cas9 mixed with crRNA and tracrRNA (Integrated DNA Technologies) and an appropriate repair template (S170A mutation: crRNA GGAGTTAGAACTCTCAGCTA, repair template TCAAATGAAGGAGTTCGAAGCTCAGAAGCACGCTATGGAAGAGCGCATTAAGGAGTTAGAGCTCGCTGCTACTGACGCTAATAATACGACCGTTGGATCATTTCGAGGAACACTCGATGATATCCTCAAGAAG; T178A mutation: crRNA GACGCTAATAATACGACCGT, repair template GAAGCACGCTATGGAAGAGCGCATTAAGGAGTTAGAACTCTCAGCTACGGACGCTAATAACACTGCAGTTGGATCATTTCGAGGAACACTCGATGATATCCTCAAGAAGAATGACCCAGACTTTACT; T198A mutation: crRNA CTGAAGTAAGAGTAAAGTCT, repair template CGGAACTCGAAGACCACATCCAACAGCTTAGACAGGAGCTCGACGACCAAGCAGCACGACTTGCAGATTCTGAAAACGTTAGAGCACAGCTGGAAGCTGCTACTGGGCAGGGTATTCTGGGA). Strains with the correct edit were identified by PCR with the following primer sets (S170A & T178A: GAAGCGTCGCAGAAGAGAGT and CTTCCAGCTGTGCTCTAACG; T198A: TGTTCAAGAGAAAATGGAGCAA and GTTTGGGACCATTGCGTTAC) followed by digestion with a diagnostic restriction site introduced during the edit (S170A: SacI; T178A: PstI; T198A: Bsu36I) and then confirmed by sequencing.

### RNA interference

Single-stranded RNAs (ssRNAs) were synthesized in 50 μL T3 and T7 reactions (MEGAscript, Invitrogen) using gel purified DNA templates generated by PCR from N2 genomic DNA using oligonucleotides containing T3 or T7 promoters (**Table S2**). Reactions were cleaned using the MEGAclear kit (Invitrogen), and the 50 μL T3 and T7 reactions were mixed with 50 μL of 3x soaking buffer (32.7 mM Na_2_HPO4, 16.5 mM KH_2_PO_4_, 6.3 mM NaCl, 14.1 mM NH_4_Cl) and annealed (68°C for 10 min followed by 37°C for 30 min). For the depletions, double-stranded RNAs (dsRNAs) were injected at a concentration of at least 1.3 μg/μL. L4 hermaphrodites were injected with dsRNA and incubated at 16°C. To assess embryonic lethality after RNAi-mediated depletion, L4 hermaphrodites were injected with dsRNA and incubated at 16°C for 48 hours. Worms were singled and allowed to lay embryos at 16°C for 24 hours. Adult worms were removed, and all embryos and hatchlings were counted after an additional 24 hours (**Fig. S1C**). For live imaging of early embryos after RNAi, L4 hermaphrodites were injected with dsRNAs and incubated at 16°C for 48 hours before dissection to isolated embryos for imaging.

### Antibodies

Antibodies against SPD-5 (392-550 aa; used at 1 μg/mL for immunoblotting; (Dammermann et al., 2004)) and γ-tubulin (428–444 aa; used at 1 μg/mL for immunoblotting; (Hannak et al., 2001)) were previously described. Antibodies against GIP-1 (used 1 μg/mL for immunoblotting) were generated by injecting a GST fusion with GIP aa 1–150 into a rabbit and affinity purifying the antibodies from serum using standard procedures (Harlow and Lane, 1988). The antibody against phosphorylated T198 in SPD-5 was raised against a phosphorylated peptide KNDPDF-[pT]-LTSGYEE, where [pT] is a phosphorylated threonine residue (Pacific Immunology). The following antibodies were purchased from commercial sources, with their working concentrations indicated in parentheses: anti-α-tubulin (1:5,000 for immunoblotting; DM1A; Sigma-Aldrich); anti-FLAG (1:1,000 for immunoblotting; F1804; Sigma-Aldrich); anti-Myc (1:5,000 for immunoblotting; monoclonal 9E10; M4439; Sigma-Aldrich). Secondary antibodies were purchased from Jackson ImmunoResearch and GE Healthcare.

### Live imaging

Embryos for live imaging experiments were obtained by dissecting gravid adult hermaphrodites in M9 buffer (42 mM Na_2_HPO4, 22 mM KH_2_PO_4_, 86 mM NaCl, and 1mM MgSO_4_). One-cell embryos were transferred with a mouth pipette onto a 2% agarose pad, overlaid with an 22 × 22-mm coverslip, and imaged using a spinning disk confocal system (Andor Revolution XD Confocal System; Andor Technology) with a confocal scanner unit (CSU-10; Yokogawa) mounted on an inverted microscope (TE2000-E; Nikon) equipped with a 60X 1.4 Plan-Apochromat objective or a 100X 1.4 Plan-Apochromat objective, solid-state 100-mW lasers, and an electron multiplication back-thinned charge-coupled device camera (iXon; Andor Technology), or an inverted microscope (Axio Observer.Z1; Carl Zeiss) equipped with a spinning-disk confocal head (CSU-X1; Yokogawa) and a 63X 1.4 NA Plan Apochromat lens (Zeiss) or a 100X 1.3 EC Plan Neofluar lens (Zeiss), in a temperature-controlled room at 20°C. Images of centrosomal fluorescence and chromosome segregation were acquired every 30 or 60s by collecting 9-12 z-planes at 1.0 µm intervals or 11 z-planes at 1.5 µm intervals without binning. Imaging was initiated in one-cell embryos between centrosome separation and pronuclear meeting and was terminated after initiation of cytokinesis.

### Image analysis

All images were processed and analyzed using ImageJ (National Institutes of Health). For figure construction, final image panels were scaled for presentation in Photoshop (Adobe Creative Cloud). A gamma of 1.5 was applied for images of mCherry and GFP-tagged SPD-5, and a gamma of 1.2 was applied to images of γ-tubulin::mCherry, GFP::MZT-1, GIP-2::GFP and GFP:: β-tubulin to allow visualization of centrosomes in the mutants, while not oversaturating the signal in the wild-type centrosomes. Centrosomal fluorescence in Figures 1B-C, 2B-G, 3I, S1F, S2C, and S3A were quantified from maximum intensity projections of entire z-stacks. Centrosomal PLK-1::GFP fluorescence (Figures 1B and S1F) and microtubule intensity (Figure 4C) were quantified from maximum intensity projections of the portion of the z-stack containing the centrosomes. To quantify centrosomal fluorescence, a fixed size box was drawn around the centrosome at each time point (smallest box that could enclose the centrosomal signal at their largest point in the image sequence; box size varied depending on marker and imaging conditions), along with a box one pixel larger on each side in both dimensions. The per-pixel background was calculated as [(integrated intensity in the larger box - integrated intensity in the smaller box)/(area of larger box - area of smaller box)]. The centrosomal signal was the integrated intensity in the smaller box minus the area of the smaller box multiplied by the per pixel background.

### Protein expression and purification

GST-tagged SPD-5 N terminus (SPD-5^N^) proteins were expressed in BL21(DE3)pLysS E. coli from DNA constructs cloned into a pGEX-6P-1 vector. When the bacterial cultures reached an OD600 of 0.6, protein expression was induced for 16-18 hours at 15°C by addition of IPTG to 0.3 mM. Cells were washed once with cold PBS and flash frozen in liquid nitrogen. Pelleted cells were resuspended in PBS with 250 mM NaCl, 10 mM EGTA, 10 mM EDTA, 0.1% Tween, 200 μg/mL lysozyme, 2 mM benzamidine, and EDTA-free protease inhibitor cocktail (Roche), and were lysed by sonication. After 20-minute centrifugation at 40,000 rpm in a 45 Ti rotor (Beckman) at 4 °C, cleared cell lysates were incubated with glutathione agarose (Sigma) for 2 hours at 4°C. The resin was then washed 2x 30 mL with washing buffer (PBS containing 250 mM NaCl, 1 mM β-mercaptoethanol, and 2 mM benzamidine) followed by incubation with 10 mL washing buffer containing 5mM ATP for 10 min at 4°C to reduce non-specific interactions with heatshock proteins. After the incubation, the resin was further washed 3 times with washing buffer and incubated with PreScission protease (Eton Bioscience) in elution buffer (20 mM Tris-Cl pH 8.0, 150 mM NaCl and 1 mM DTT) overnight at 4°C to elute the SPD-5^N^ protein by cleavage from the GST-tag. The supernatant was collected the next day and stored at 4°C. PLK-1 T194D, purified from Sf9 cells, was a gift from Jeffrey Woodruff (UT Southwestern).

### Purification of reconstituted *C. elegans* γ-tubulin complexes

The *C. elegans* γ-tubulin complex was reconstituted by co-expression in human FreeStyle 293-F cells (Thermo Fisher Scientific). The GIP-2 and γ-tubulin (TBG-1) coding sequences were amplified by PCR from an N2 cDNA library. For MZT-1 and GIP-1, coding sequences optimized for human cell expression were synthesized (GENEWIZ). Sequences encoding Myc-tagged GIP-2, FLAG-tagged MZT-1, GIP-1 or *C. elegans* γ-tubulin were cloned into human expression vectors driven by the CMV promoter (Table 3) and co-transfected into FreeStyle 293-F cells (Thermo Fisher Scientific). The empty 5Myc plasmid (CS2P, Addgene #17095) or 3FLAG plasmid (p3XFLAG-CMV^™^-7.1, SIGMA #E7533) were used as negative controls. Cell transfection was performed using FreeStyle MAX Reagent and OptiPRO SFM according to the manufacturer’s guidelines (Thermo Fisher Scientific). 10 mL of cells at 1⨯ 10^6^ cells/mL were transfected with a total of 12.5 µg DNA constructs. 43-48 hours after transfection, cells were harvested and washed with PBS. The cells were resuspended in lysis buffer (20 mM Tris/HCl pH 7.5, 50 mM NaCl, 1% Triton X-100, 5 mM EGTA, 1 mM dithiothreitol (DTT), 2 mM MgCl_2_ and EDTA-free protease inhibitor cocktail (Roche)) and lysed in an ice-cold sonicating water bath for 5 minutes. After 15-minute centrifugation at 15,000x g. at 4 °C, the whole cell lysates were incubated with Pierce Anti-c-Myc magnetic beads (Thermo Fisher Scientific) for 2 hours at 4 °C. The beads were washed five times with lysis buffer and used for pull-down assay or resuspended in SDS sample buffer. For immunoblotting, equal volumes of samples were run on Mini-PROTEAN gels (Bio-Rad) and transferred to PVDF membranes using a TransBlot Turbo system (Bio-Rad). Blocking and antibody incubations were performed in TBS-T plus 5% nonfat dry milk or in TBS-T plus 5% BSA. Immunoblotting was performed as described above.

### Kinase and Pulldown assays

For the pulldown assays in Figure 3, 1.5μM SPD-5^N^ proteins were mixed with 200 nM constitutively active PLK-1 T194D (gift from Jeffrey Woodruff) in kinase buffer (20 mM Tris-Cl pH 7.5, 50 mM NaCl, 10 mM MgCl2, 0.2 mM ATP, 1 mM DTT). After incubation for 1 hour at 23°C, proteins were mixed with Myc beads bound γ-tubulin complexes in lysis buffer and incubated for 2 hours at 4°C. The final concentration of SPD-5^N^ was 75nM. The beads were washed five times with lysis buffer and resuspended in sample buffer before analysis on SDS-PAGE.

### Screen for PLK-1 putative phosphorylation sites

Potential PLK-1 phosphorylation sites in SPD-5 were identified using the kinase-specific phosphorylation site prediction system (http://polo.biocuckoo.org/down.php). Protein sequences in FASTA format were entered and threshold setting was set to ALL. A list of candidate PLK-1 sites for each target protein was generated utilizing the algorithm shown in Figure S1D and the selected sites were mutated in regional clusters.

---

**Table S1.** *C. elegans* strains used in this study.

| Strain # | Genotype | Figure |
| --- | --- | --- |
| N2 | wild type (ancestral) | 1D, 2A |
| OD868 | ItSi220[pOD1249/pSW077; Pmex-5::GFP-tbb-2-operon-linker-mCherry-his-11; cb-unc-119(+)]I | 1D, 2I |
| OD1709 | ItSi569[oxTi185; pOD1110/pSW008; CEOP3608 TBG-1::mCherry; cb-unc-119(+)]I; unc-119(ed3)III | 3E, S2D |
| OD1801 | ItSi592[pVV168; Pspd-2::GFP-spd-5 mut 653,658, reencoded; cb-unc-119(+)]II; unc-119(ed3)III | 2A |
| OD2283 | ItSi569[oxTi185; pOD1110/pSW008; CEOP3608 TBG-1::mCherry; cb-unc-119(+)]I; ItSi592[pVV168; Pspd-2::GFP-spd-5 mut 653,658, reencoded; cb-unc-119(+)]II; unc-119(ed3)III | 1C, 2D, S2D |
| OD2433 | ItSi1141[pOD1021/pVV103; Pspd-2::GFP::SPD-5 reencoded; cb-unc-119(+)]II; unc-119(ed3) III | 2A |
| OD2435 | ItSi569[oxTi185; pOD1110/pSW008; CEOP3608 TBG-1::mCherry; cb-unc-119(+)]I; ItSi1141[pOD1021/pVV103; Pspd-2::GFP::SPD-5 reencoded; cb-unc-119(+)]II; unc-119(ed3) III | 1C, 2B, 2H, S2D |
| OD2887 | ItSi937[pOD2079/pSW400; Pspd-2::GFP::SPD-5(T335A, S339A, S340A, S387A); cb-unc-119(+)]II; unc-119(ed3)III | 2A |
| OD2888 | ItSi938[pOD2080/pSW401; Pspd-2::GFP::SPD-5(S441A, S475A, S479A, T488A); cb-unc-119(+)]II; unc-119(ed3)III | 2A |
| OD2889 | ItSi939[pOD2081/pSW402; Pspd-2::GFP::SPD-5(S558A, S627A, S635A, S649A); cb-unc-119(+)]II; unc-119(ed3)III | 2A |
| OD2890 | ItSi940[pOD2082/pSW403; Pspd-2::GFP::SPD-5(S688A, T755A, T1070A); cb-unc-119(+)]II; unc-119(ed3)III | 2A |
| OD3636 | ItSi569[oxTi185; pOD1110/pSW008; CEOP3608 TBG-1::mCherry; cb-unc-119(+)]I; ItSi1560[pOD1021/pVV103; Pspd-2::GFP::SPD-5 T178A reencoded; cb-unc-119(+)]II; unc-119(ed3) III. Note WT transgene was in vivo CRISPR edited to introduce the T178A mutation. | 3E |
| OD3760 | ItSi569[oxTi185; pOD1110/pSW008; CEOP3608 TBG-1::mCherry; cb-unc-119(+)]I; ItSi1556[pOD1021/pVV103; Pspd-2::GFP::SPD-5 T198A reencoded; cb-unc-119(+)]II; unc-119(ed3) III. Note WT transgene was in vivo CRISPR edited to introduce the T198A mutation. | 3E, S2D |
| OD3778 | ItSi569[oxTi185; pOD1110/pSW008; CEOP3608 TBG-1::mCherry; cb-unc-119(+)]I; ItSi1558[pOD1021/pVV103; Pspd-2::GFP::SPD-5 S170A reencoded; cb-unc-119(+)]II; unc-119(ed3) III. Note WT transgene was in vivo CRISPR edited to introduce the S170A mutation. | 3E |
| OD4047 | unc-119(ed3)III; ItSi1553[pMO86; Pspd-2::spd-2 reencoded::spd-2 3'UTR; cb-unc119(+)]V | 1D |
| OD4049 | unc-119(ed3)III; ItSi1555[pMO87; Pspd-2::spd-2 S232T, T233A reencoded::spd-2 3'UTR; cb-unc119(+)]V | 1D |
| OD4098 | spd-5(It102[mcherry::spd-5])I; plk-1(It18[plk-1::sGFP]:loxp)III; unc-119(ed3)III; ItSi1553[pMO86; Pspd-2::spd-2 reencoded::spd-2 3'UTR; cb-unc119(+)]V | 1B |
| OD4099 | spd-5(It102[mcherry::spd-5])I; plk-1(It18[plk-1::sGFP]:loxp)III; unc-119(ed3)III; ItSi1554[pMO87; Pspd-2::spd-2 S232T, T233A reencoded::spd-2 3'UTR; cb-unc119(+)]V | 1B |
| OD4166 | ItSi220[pOD1249/pSW077; Pmex-5::GFP-tbb-2-operon-linker-mCherry-his-11; cb-unc-119(+)]I unc-119(ed3)III; ItSi1554[pMO87; Pspd-2::spd-2 S232T, T233A reencoded::spd-2 3'UTR; cb-unc119(+)]V | 1D, 4A, 4C |
| OD4168 | ltSi220[pOD1249/pSW077; Pmex-5::GFP-tbb-2-operon-linker-mCherry-his-11; cb-unc-119(+)]I unc-119(ed3)III; ltSi1552[pMO86; Pspd-2::spd-2 reencoded::spd-2 3'UTR; cb-unc119(+)]V | 1D, 4A<br>4C |
| OD4181 | ltSi1532[pMO91; Pspd-2::gfp::spd-5 S5A,S11A::spd-5 3'UTR; cb-unc119(+)]II; unc-119(ed3) III | 2A |
| OD4227 | gip-2(lt19[gip-2::GFP]::loxP::cb-unc-119(+)::loxP)I; sas-4(lt128[mcherry::sas-4])III; ltSi1129[pZZ2; Pspd-2::SPD-5 (re-encoded);cb-unc-119(+)]II; unc-119(ed3)III | S3A, S3B |
| OD4235 | ltSi220[pOD1249/pSW077; Pmex-5::GFP-tbb-2-operon-linker-mCherry-his-11; cb-unc-119(+)]I; ltSi1129[pZZ2; Pspd-2::SPD-5 (re-encoded) (WT, untagged);cb-unc-119(+)]II; unc-119(ed3)III | 1D, 2I,<br>4B, 4C |
| OD4313 | ltSi220[pOD1249/pSW077; Pmex-5::GFP-tbb-2-operon-linker-mCherry-his-11; cb-unc-119(+)]I; ltSi1219[pMO104; Pspd-2::spd-5 S653A S658A::spd-5 3'UTR; cb-unc119(+)]II; unc-119(ed3)III | 1D, 2I<br>4B, 4C |
| OD4465 | ltSi1539[pMO130; Pspd-2-GFP::spd-5 S170A, T178A, T198A::spd-5 3'UTR; cb-unc119(+)]II; unc-119(ed3)III | 2A |
| OD4467 | ltSi220[pOD1249/pSW077; Pmex-5::GFP-tbb-2-operon-linker-mCherry-his-11; cb-unc-119(+)]I; ltSi1232 [pMO123; Pspd-2::spd-5 S170A, T178A, T198A::spd-5 3'UTR; cb-unc119(+)]II; unc-119(ed3)III | 2I, 4B, 4C |
| OD4495 | ltSi569[oxTi185; pOD1110/pSW008; CEOP3608 TBG-1::mCherry; cb-unc-119(+)]I; ltSi1539[pMO130; Pspd-2-GFP::spd-5 S170A, T178A, T198A::spd-5 3'UTR; cb-unc119(+)]II; unc-119(ed3)III | 2C, 2H,<br>S2D |
| OD4680 | gip-2(lt19[gip-2::GFP]::loxP::cb-unc-119(+)::loxP)I; ltSi1232 [pMO123; Pspd-2::spd-5 S170A, T178A, T198A::spd-5 3'UTR; cb-unc119(+)]II; unc-119(ed3)III; sas-4(lt128[mcherry::sas-4])III | S3A, S3B |
| OD4746 | ltSi1523[pMO172; Pspd-2::gfp::spd-5 S240A,T274A::spd-5 3'UTR; cb-unc119(+)]II;unc-119(ed3)III; | 2A |
| OD4828 | spd-2(lt143[mCherry::spd-2]) I; ltSi1129[pZZ2; Pspd-2::SPD-5 (re-encoded);cb-unc-119(+)]II; plk-1(lt18[plk-1::sGFP]::loxP)III; unc-119(ed3)III | S1F |
| OD4829 | spd-2(lt143[mCherry::spd-2]) I; ltSi1232 [pMO123; Pspd-2::spd-5 S170A, T178A, T198A::spd-5 3'UTR; cb-unc119(+)]II; ; plk-1(lt18[plk-1::sGFP]::loxP)III; unc-119(ed3)III | S1F |
| OD4833 | ltSi220[pOD1249/pSW077; Pmex-5::GFP-tbb-2-operon-linker-mCherry-his-11; cb-unc-119(+)]I; ltSi1561 [pMO174; Pspd-2::spd-5 S170A, T178A, T198A, S653A, S658A::spd-5 3'UTR; cb-unc119(+)]II; unc-119(ed3)III | 4B, 4C |
| OD4840 | ltSi569[oxTi185; pOD1110/pSW008; CEOP3608 TBG-1::mCherry; cb-unc-119(+)]I; ltSi1129[pZZ2; Pspd-2::SPD-5 (re-encoded);cb-unc-119(+)]II; unc-119(ed3)III | 2E, 3H,<br>3I, S2D |
| OD4841 | ltSi569[oxTi185; pOD1110/pSW008; CEOP3608 TBG-1::mCherry; cb-unc-119(+)]I; ltSi1218 [pMO104; Pspd-2::spd-5 S653A S658A::spd-5 3'UTR; cb-unc119(+)]II; unc-119(ed3)III | 2G, S2D |
| OD4843 | ltSi569[oxTi185; pOD1110/pSW008; CEOP3608 TBG-1::mCherry; cb-unc-119(+)]I; ltSi1232 [pMO123; Pspd-2::spd-5 S170A, T178A, T198A::spd-5 3'UTR; cb-unc119(+)]II; unc-119(ed3)III | 2F, 3I,<br>S2D |
| OD4870 | mzt-1(wow51[gfp::mzt-1]) I; ltSi1129[pZZ2; Pspd-2::SPD-5 (re-encoded); cb-unc-119(+)]II; unc-119(ed3)III | S2B, S2C |
| OD4879 | ltSi569[oxTi185; pOD1110/pSW008; CEOP3608 TBG-1::mCherry; cb-unc-119(+)]I; ltSi1214[pMO095; Pspd-2::spd-5 T198A (re-encoded)::spd-5 3'UTR; cb-unc119(+)]II; unc-119(ed3)III | 3H, 3I,<br>S2D |
| OD4880 | mzt-1(wow51[gfp::mzt-1]) I; ItSi1214[pMO095; Pspd-2::spd-5 T198A (re-encoded)::spd-5 3'UTR; cb-unc119(+)]II; unc-119(ed3)III | S2B, S2C |
| OD4882 | mzt-1(wow51[gfp::mzt-1]) I; ItSi1232 [pMO123; Pspd-2::spd-5 S170A, T178A, T198A::spd-5 3'UTR; cb-unc119(+)]II; unc-119(ed3)III | S2B, S2C |

**Table S2.**
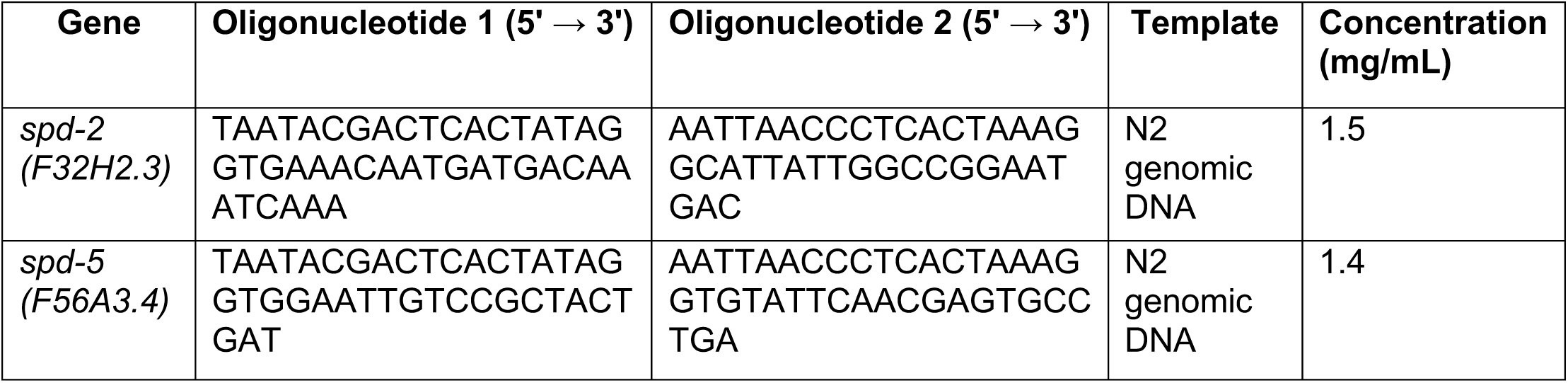
Oligos used for dsRNA production.

**Table S3.** Plasmids used in this study.

| plasmid # | Description | Bacterial selection | Figure |
| --- | --- | --- | --- |
| **pOD3898** | pCMV-TBG-1 (*C. elegans* γ-tubulin) | Ampicillin | 3A, 3C, 3D, 3G |
| **pOD3899** | pCMV-GIP-2-5Myc | Ampicillin | 3A, 3C, 3D, 3G |
| **pOD3900** | pCMV-GIP-1 | Ampicillin | 3A, 3C, 3D, 3G |
| **pOD3901** | pCMV-3FLAG-MZT-1 | Ampicillin | 3A, 3C, 3D, 3G |
| **pOD3902** | pGEX-6P-1-GST-SPD-5 (aa1-473) | Ampicillin | 3B, 3C, 3D, 3F, 3G, S2A |
| **pOD3903** | pGEX-6P-1-GST-SPD-5 (aa1-473) T198A | Ampicillin | 3F, 3G, S2A |
| **pOD3904** | pGEX-6P-1-GST-SPD-5 (aa1-473) S170A, T178A, T198A | Ampicillin | 3B, 3C, 3G, S2A |

